# Population size shapes the evolution of lifespan

**DOI:** 10.1101/2022.12.17.520867

**Authors:** Martin Bagic, Dario Riccardo Valenzano

## Abstract

Biological aging results from the age-dependent change in the force of natural selection, which increases the probability of germline variants that limit survival to accumulate in genes acting predominantly in late life^1^. The evolutionary mechanisms underlying the accumulation of neutral mutations and antagonistically pleiotropic gene variants that cause biological aging have been analyzed to date under the assumption of infinitely large population size. However, even though population size importantly shapes genetic and phenotype variation via drift and selection^2,3^, we still have a limited understanding of how finite population size impacts the evolution of mortality at the population level. Here, we study the impact of population size on lifespan evolution under mutation accumulation and antagonistic pleiotropy. We found that larger population size leads to lower age-independent, as well as age-dependent mortality under mutation accumulation, due to more effective purifying selection against deleterious germline variants. Strikingly, large population size can lead to extended lifespan under antagonistic pleiotropy, due to more effective positive selection on gene variants increasing survival in early-life, while leading to increased post-maturation mortality. Our findings provide a comprehensive numerical framework for the two major evolutionary genetic theories of aging and reveal a fundamental and yet non-appreciated role for population size in the evolution of mortality trajectories.

## Main

The changing force of natural selection as a function of age has been long identified as a leading cause for the accumulation of deleterious gene variants that impact organism survival and aging^4,5^. Indeed, as soon as individuals are able reproduce, deleterious gene variants affecting late-life phenotypes are transmitted to the next generation even before their phenotypic effect becomes selectable. As a result, deleterious germline mutations tend to accumulate in genes that contribute to late-life maintenance and survival, a phenomenon labeled aging by “mutation accumulation”^1^. Additionally, aging-causing variants can also accumulate as positively selected alleles, whenever an overall selectively advantageous variant negatively impacts survival at advanced ages – a phenomenon labeled aging by antagonistic pleiotropy^6^. Hence, while the mutation accumulation theory of aging^4^ is compatible with the accumulation of neutral to nearly neutral variants in genes affecting late life phenotypes, the antagonistic pleiotropy theory of aging is consistent with the evolution of gene variants under positive directional selection^6^.

Mutation-selection balance importantly affects the accumulation of mutations with age-specific effects^7,8^, e.g. when the selective advantage of a given mutation becomes more prominent at advanced ages, rather than in early life. However, the context under which selection and drift can impact the accumulation of aging-relevant mutations has to date been surprisingly poorly explored. In this work, we aim to fill this important gap in knowledge. The analysis of the evolution of senescence has been largely conducted under the assumption of infinitely large population size. However, population size importantly affects the probability of fixation of beneficial, as well as nearly neutral variants^1,3,9^. While large population size leads to more effective fixation probability for beneficial variants, small population size can lead to increased chances for neutral and slightly deleterious mutations to reach high frequencies and even to reach fixation^10,11^. Since aging-determining variants can accumulate under both drift and selection, here we asked what the impact of population size is on the molding of senescence.

### Evolving populations under mutation accumulation and antagonistic pleiotropy

We developed a numerical model (AEGIS^12^ and **Methods**) to simulate the evolution of gene variants affecting survival and reproduction at discrete ages, with or without pleiotropic benefits on individual survival (**Figure 1**). We let populations of independent agents that reproduce either sexually or asexually, to evolve age-specific survival probabilities under mutation, selection and drift in populations of different sizes. While deleterious mutations affected individual, age-specific survivorship in the mutation-accumulation framework, we enabled antagonistic pleiotropy through matrix multiplication, where individual gene variants affect survival at two different time points, with the rule that survival is increased at the earlier time point and lowered at the later time point (**Figure 1** and **Methods**). Following this setup, we let populations evolve age-specific mortality in a timeframe of up to 10^5^ overlapping generations under limiting resources (**Methods**). We ran a total of 360 simulations across different population sizes (N= 3x10^2^, 3x10^3^ and 3x10^4^), reproductive mode (sexual and asexual reproduction), mutation accumulation and antagonistic pleiotropy (**Methods**). Each simulation was initialized under a “non-aging” condition, consisting of constant, age-specific mortality (5% per time stage). Hence, during each simulation, changes in age-dependent mortality evolve as the outcome of the accumulation of beneficial or detrimental variants under drift and selection in gene bits affecting age-specific mortality (**Methods**).

**Figure 1.**
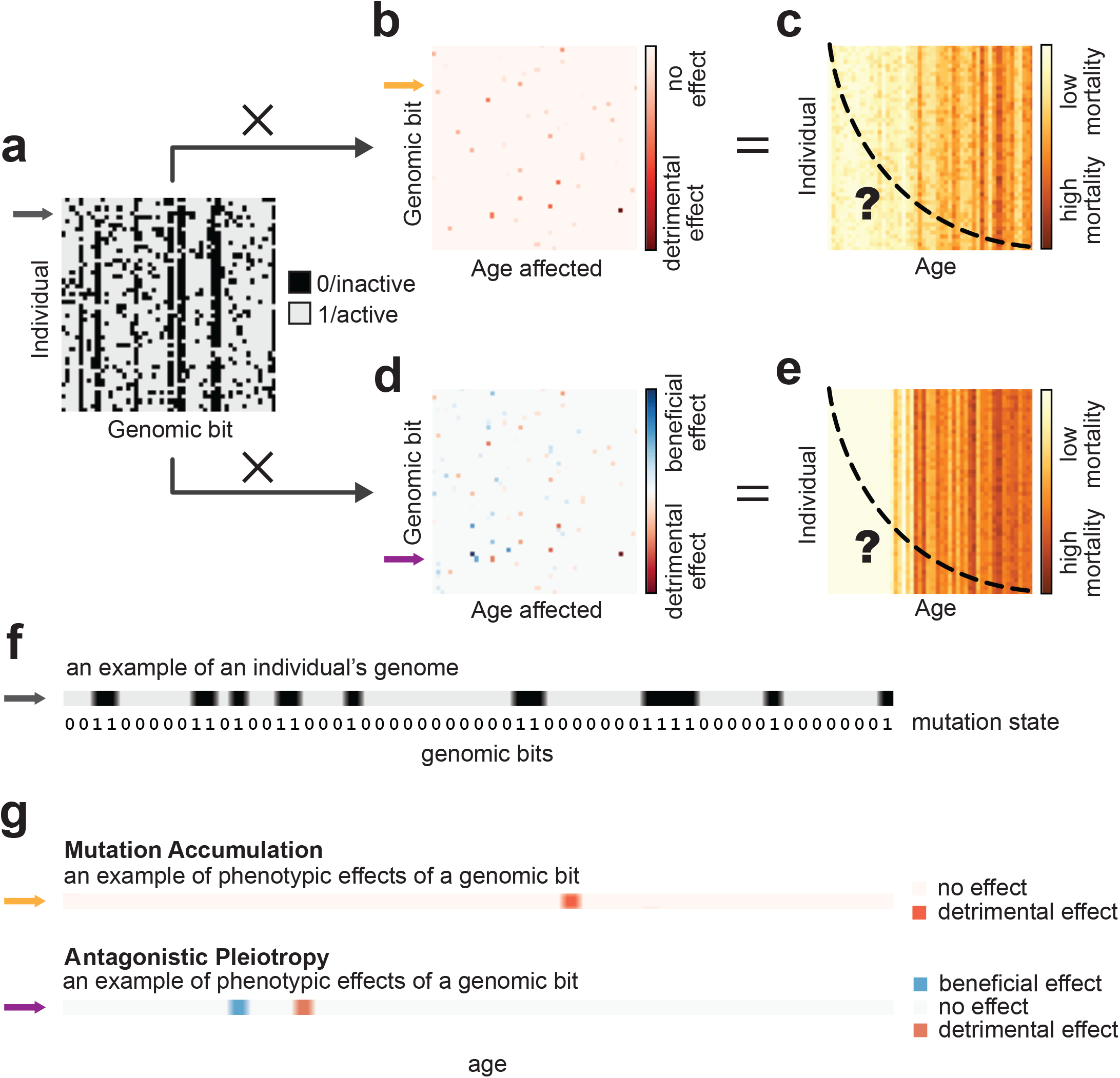
Genotype-to-phenotype mapping. Every individual carries a genomic bitstring composed of genomic bits which can be in the inactive or the active state (**a**, an example in **f**). Mortality phenotypes (**c**,**e**) are computed by multiplying the genomic bitstrings (**a**) with genotype-to-phenotype matrices (**b**,**d**). Under mutation accumulation, every genomic bit has an age-specific detrimental effect (**b**, an example in **g**), while under antagonistic pleiotropy they also have a beneficial effect at an earlier age (**d**, an example in **g**). Genotype-to-phenotype matrices (**b**,**d**) are defined before the start of the simulation while genomic bitstrings (**a**), mortalities and survival curves (**c**,**e**) evolve during the simulation.

### Larger population size mitigates the load of deleterious mutations in early life under mutation accumulation

Mutation load increases in genes preferentially acting in late-life, due to their decreased gene-specific effective population size; a phenomenon known as “selection shadow”^13-15^.

We asked whether under no pleiotropy, i.e. in a purely “mutation accumulation” framework, population size affected the evolution of age-specific mortality (**Figure 2** and **Extended Data Figure 1**). We found that population size affected mortality at all ages. Larger population size led to lower mortality across all ages, consistent with lower genome-wide mutation load in larger populations.

**Figure 2.**
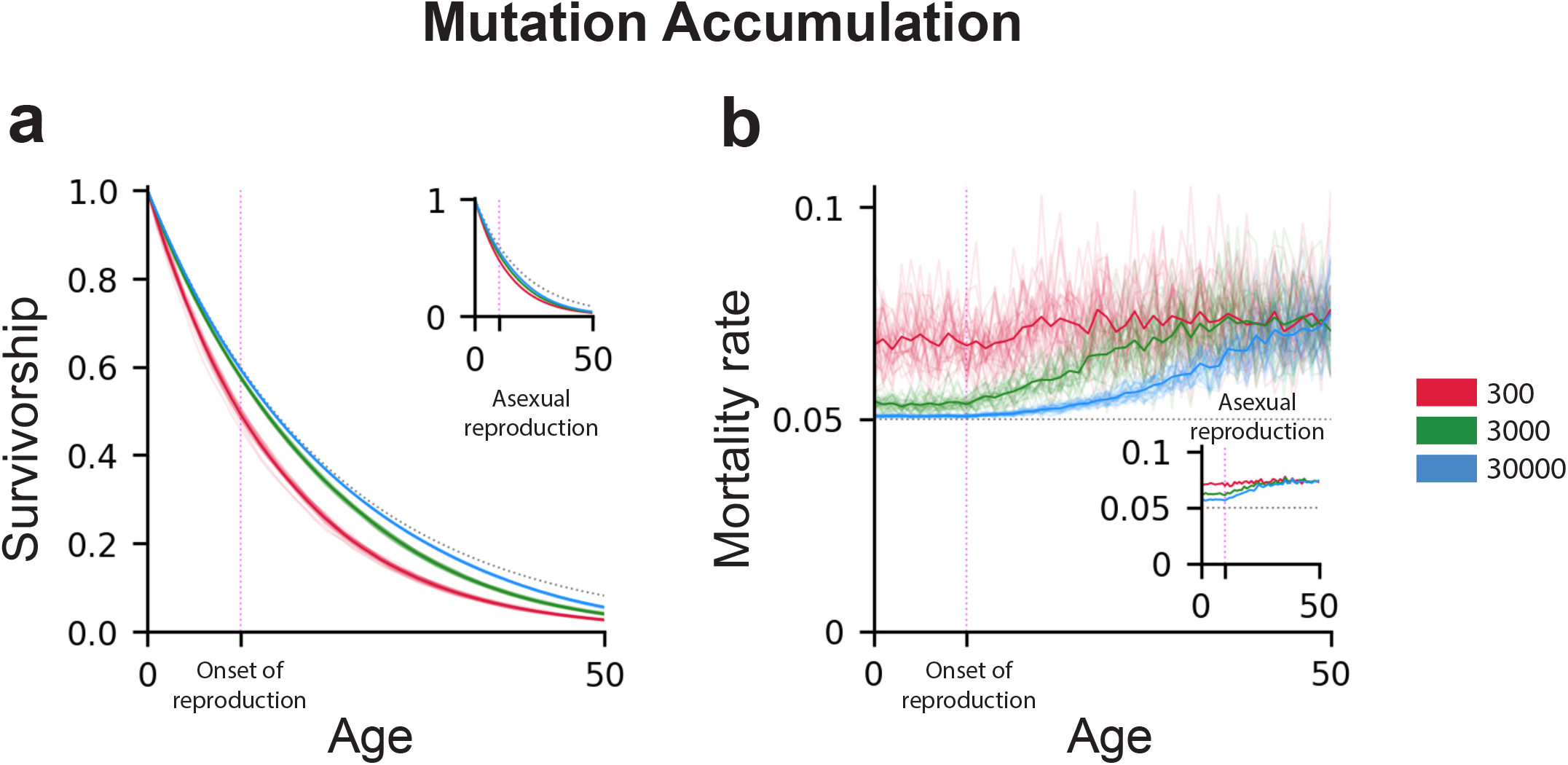
Population size impacts the evolution of mutations deleterious in early life. Populations of different sizes evolve different survival (**a**) and mortality (**b**) curves, when evolving under mutation accumulation. Initially, all populations are phenotypically identical (gray dotted curves) but they differ at the end of the simulation (thin colored curves show the phenotypes of individual simulations; the thick colored curves show the mean phenotypic curve). By selecting against deleterious mutations more efficiently, larger populations retain lower mortality in early life (**b**) which only begins to increase at a later age, after the onset of reproduction. The same is true for asexually reproducing populations; however, the mortalities in such populations are higher and they start to increase earlier than in sexually reproducing populations.

In simulations with larger population sizes, we found that genomes accumulated fewer deleterious variants in genes that affect early survival compared to genes that affect late survival.

Despite living longer, larger populations evolve a more rapid increase in mortality trajectories than smaller populations, due to stronger purifying selection in early life (**Figure 2b**). Hence, larger populations evolve to be longer lived and have late-life compressed mortality compared to smaller populations.

We ran simulations in populations reproducing either sexually or asexually (**Methods**). Overall, we found that irrespective of the reproductive modality, larger populations led to lower mortality. However, given same population size, sexual reproduction was more effective in purging genomes from survival-detrimental variants, resulting in lower mortality compared to the asexual model (**Figure 2**, insets).

### Under antagonistic pleiotropy, larger population size leads to longer life and compressed late-life mortality

We allowed populations to evolve genes that, while promoting survival in early life, caused a survival penalty in late life (**Figure 1, Extended Data Figure 1** and **Methods**). Under this scenario, compared to the initial state consisting in constant mortality across all ages, positive selection for beneficial variants led populations to evolve extended life in both small and large populations (**Figure 3a**). However, in larger populations adult mortality accelerated later than in smaller populations (**Figure 3b**). Furthermore, under larger population sizes, adult mortality became compressed, resulting in the rectangularization of survival^16^. Hence, larger populations evolved to be longer-living (delayed aging) while having higher late-life mortality rates (faster aging) than smaller populations.

**Figure 3.**
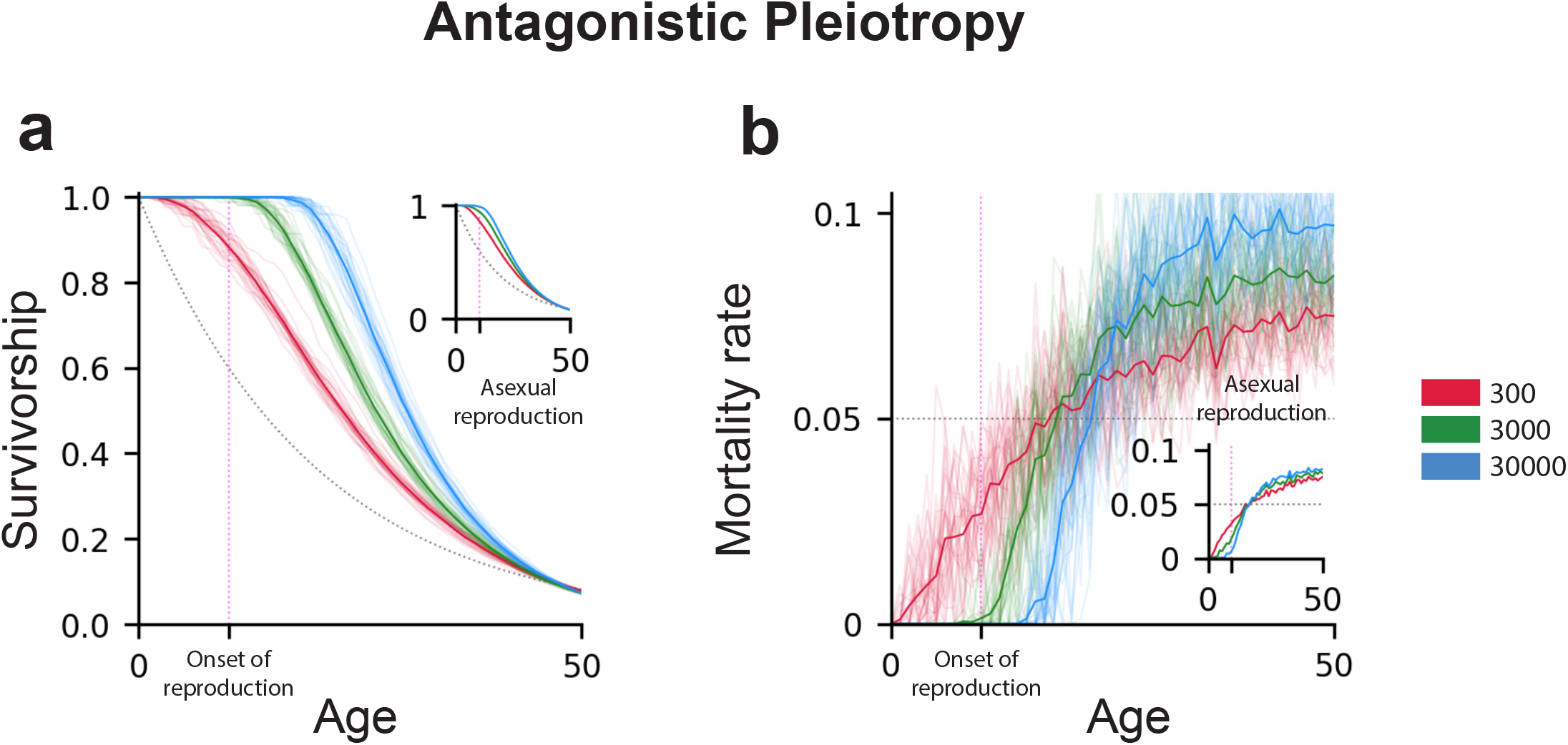
Population size impacts the evolution of pleiotropic variants. Populations of different sizes evolve different survival (**a**) and mortality (**b**) curves, when evolving under mutation accumulation. Initially, all populations are phenotypically identical (gray dotted curves) but they differ at the end of the simulation (thin colored curves show the phenotypes of individual simulations; the thick colored curves show the mean phenotypic curve). By selecting for fitness-increasing mutations more efficiently, larger populations evolve lower early-life mortality, extending lifespan (**a**) and delaying the onset of aging (**b**). However, due to the more intense accumulation of such pleiotropic mutations, larger populations evolve higher late-life mortality (**b**) resulting in a steeper decline in survivorship (**a**) and more rapid aging.

We evolved antagonistic pleiotropy under both sexual and asexual reproduction (**Methods** and **Figure 3**, insets). Irrespective of the reproductive modality, larger population size resulted in the evolution of longer life. However, sexual reproduction resulted in more effective fixation of beneficial variants that led to higher early life survival and to faster late-life mortality acceleration compared to the asexual model (**Figure 3**, insets).

### Lifespan variation evolves as a function of population size

To explicitly answer whether changes in population size led to changes in population median lifespan, we ran a total of 180 simulations varying 15 population sizes, ranging from 100 to 3000. We found that both under the sexual and the asexual model, median lifespan evolved as a function of population size, with large populations evolving longer median lifespan under both mutation accumulation and antagonistic pleiotropy (**Figure 4** and **Extended Data Figure 2**). Our results generalize empirical findings obtained in Daphnia^17^ and annual killifish^15,18^, according to which population size impacts evolution of lifespan.

**Figure 4.**
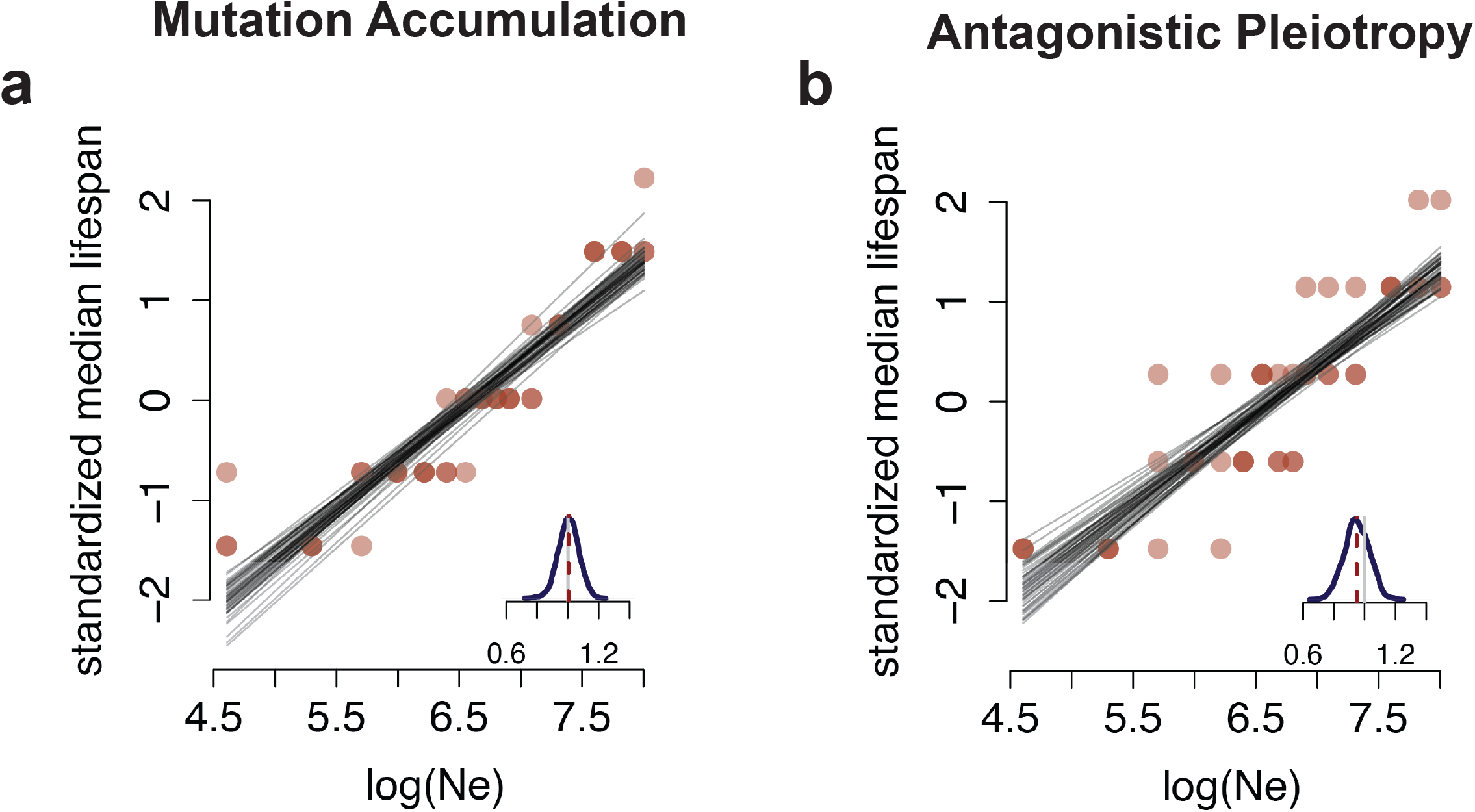
Population size impacts the evolution of lifespan. Correlation between standardized median lifespan (y-axis) and population size (log-scale, x-axis) under both mutation accumulation and antagonistic pleiotropy. Both distributions are from the “sexual reproduction” model. Each plot displays 50 regression lines (gray) sampled from the posterior distribution of each regression model. Insets: distribution of regression coefficients (slope); the vertical gray line corresponds to slope = 1 and the vertical dashed red line represents the mean slope for each model.

## Summary

At each new generation, novel germline genetic variants are introduced into the gene pool. The likelihood of these variants being passed on to future generations depends on both their selective advantage and random chance. The effective population size can impact how the frequency of each variant changes over time. A larger effective population size can lead to more effective purifying and positive selection, which can help remove deleterious mutations and increase the frequency of advantageous variants, respectively^9,19^. Moreover, larger effective population size mitigates genetic drift, while smaller effective population size makes drift a major force causing allele frequency changes over generations. Two main evolutionary forces influence whether novel genetic variants that impact aging and lifespan diffuse in population over generations. These forces are i) the waning strength of natural selection as a function of organismal age and ii) the selective advantage of variants that, despite being overall beneficial, decrease late-life survival or reproduction^1^. Since population size impacts whether deleterious and beneficial mutations are maintained or removed from the gene pool, we asked whether varying effective population size can impact the evolution of aging under both mutation accumulation and antagonistic pleiotropy. We simulated the evolution of biological aging under various population sizes and found that under both mutation accumulation and antagonistic pleiotropy, a smaller effective population size leads to the evolution of a shorter median lifespan. Under mutation accumulation, both in large and small populations, mutations negatively affecting survival in early life are more efficiently removed by purifying selection compared to those impacting late life. However, compared to larger populations, smaller populations accumulate also more mutations that reduce survival across all ages, including in early life. Hence, our findings indicate that in species with larger effective population size, the negative effects of mutation accumulation on survival are mitigated compared to smaller populations.

Under antagonistic pleiotropy, we discovered that in larger populations, individuals evolved a longer lifespan and a delayed onset of adult mortality. However, late-life mortality in larger populations becomes higher compared to smaller populations, due to the more effective selection of gene variants that have negative pleiotropic effects on survival in later life. Therefore, we discovered that in the case of antagonistic pleiotropy, smaller population size resulted in less efficient fixation of overall beneficial variants, indicating that the phenomenon of “selection shadow” also applies to antagonistic pleiotropy and is not exclusive to maladaptive mutations.

Our findings provide new evidence that demography plays a key role in the evolution of aging trajectories across populations under both mutation accumulation and antagonistic pleiotropy^20^.

## Methods

To generate the results we use AEGIS – a numerical model for simulating life history evolution in age-structured populations (<u>aegis-sim</u> v2.1; https://github.com/valenzano-lab/aegis). We study populations of various sizes (300, 3000 and 30000), reproducing sexually or asexually, and evolving under the MA or the AP regime. For each experimental setup, we run 30 simulations. We have run 360 simulations in total.

### Simulation steps

The simulation consists of 10^6 discrete stages. This amounts to approximately 10^5 generations. During each stage, three steps occur: overpopulation control, intrinsic mortality and reproduction.

#### Overpopulation control (step 1)

If the population grows beyond the user-defined maximum population size (e.g., 300, 3000 or 30000), 5% of the population dies from overcrowding. If overcrowding continues into the next stage, an even greater share of the population dies. The proportion of dying individuals is calculated as 1 − 0.95^*t*^ where *t* is the number of uninterrupted stages of overcrowding causing population size oscillations (**Extended Data Figure 3**). By culling 5% of the population upon overcrowding, AEGIS provides an age-independent, extrinsic source of mortality.

#### Intrinsic mortality (step 2)

Each individual suffers from an intrinsic probability of dying that depends on the individual-specific sets of accumulated mutations. At the beginning of the simulation every individual has a 5% chance of dying at every stage. The age-specific probability of dying can change to higher or lower rates as populations evolve.

#### Reproduction (step 3)

Every sexually mature individual (having survived 10 or more stages) has 30% chance of producing a single offspring at every stage. Note that stages are unitless (they do not specifically represent years, months, etc.). The newborns are added directly into the population; thus, the generations are overlapping (multiple generations are alive at the same time). Also, due to the nature of the numerical simulation, generations are discrete (reproduction occurs at fixed time intervals).

#### Initialization

The simulation is initialized with a population of genetically identical individuals with an intrinsic mortality rate of 5% per stage. Since initially survival does not decrease with age, the individuals in the initialized population do not exhibit aging. Therefore, any observed age-specific change in survival pattern during the simulation is the result of evolution by selection and drift. Importantly, the initial total mortality rate is higher than 5% since the individuals are not only subject to the intrinsic mortality rate (5%) but also the extrinsic mortality rate due to overcrowding. All individuals in the initial population are of age 0.

### Genetic architecture

Genomic bitstrings are arrays of 5000 bits that can be in either of the two states — 0 or 1. Mutations, then, are equivalent to a bit switching their state (from 0 to 1 or from 1 to 0). Initially, genomic bitstrings contain only 0’s (no 1’s).

#### Mutation rate

The probability of a bit 0 switching to 1 is *p*_0→1_ and the bit 1 switching to 0 is *p*_1→0_. The ratio of these probabilities is set to 100:1. The total mutation probability (*p*_1→0_ +*p*_0→1_) is set to 10^−5^.

Note that there are no “somatic mutations” accumulating over the lifetime of individuals; instead, all mutations are either inherited or emerge in the germline, thus they are carried by individuals from birth until death and they are passed onto their offspring during reproduction.

#### Recombination

In asexually reproducing populations, the offspring genomic bitstring is derived directly from its parent, while in sexually reproducing populations it is computed by joining two recombined haploid genomic bitstrings of the two parents. All alleles have additive genetic effects. In the current setup, the recombination rate is set to 50%. The higher the recombination rate, the higher the selection efficiency.

#### Mutation effects

Mutation effects are additive (there is no interaction between mutations), age-specific (mutations affect survival at a single age) and transient (subsequent ages are not affected).

We study two mutational landscapes as described by the *mutation accumulation* (MA) theory and the *antagonistic pleiotropic* (AP) theory. Under MA, mutations decrease survival; under AP, they increase survival at one age but decrease it at another, later age.

The effect sizes of mutations are exponentially distributed (Eyre-Walker and Keightley, 2007) with parameter of distribution set to 0.0007. This means that the average mutation alters survival by 0.07 percentage points (effects are additive, not multiplicative); however, most mutations have smaller effect sizes while a few mutations have much larger effect sizes.

The timings of detrimental effects 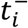(i.e., the age at which a mutation reduces survival) are uniformly distributed over the possible lifespan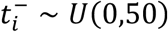. For pleiotropic mutations, the accompanying beneficial effects are uniformly distributed across ages before the detrimental effect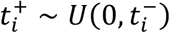.

#### Phenotype landscape

Since genomic bitstrings are diploid, every bit has a counterpart with identical mutation effects; thus, the genomic bitstring contains 2500 pairs of bits.

The number of different life histories that the simulation can produce is limited since the mutation effects for each bit are determined during the initialization phase and do not change during the simulation. Given 2500 pairs of bits, the simulation can produce up to 3^2500 or roughly 10^1193 different phenotypes.

### Genotype-to-phenotype computation

Phenotype P can be computed from a genotype *G* by a vector multiplication GM+0.95 = P where **M** is the matrix containing mutation effects (**Figure 1**).

G is a row vector of size 5000 (containing 5000 bits) and phenotype P is a row vector of size 50 (containing 50 float values encoding the age-dependent probability of survival).

M is a matrix of dimension 5000x50. Each row in M encodes all the phenotypic effects of one genomic bit. Under the MA regime, all cells in a single row should be 0 except one, which should be less than 0. On the other hand, under the AP regime, only two cells should be non-zero – one containing a negative and another one a positive value.

To compute phenotypes of the whole population, a matrix of genotypes composed of individual row genotype vectors (**G**) is multiplied with the matrix containing mutational effects (**M**) to yield a matrix of corresponding phenotypes (**P**).

### Simulation time vs generation time

Each simulation consists of 10^6 stages (each consisting of an *overpopulation control* step, *intrinsic mortality* step and *reproduction* step). The maximum number of generations is 10^5 since age of maturity is 10 (10^6/10=10^5); however, the true number of generations is lower because many individuals are born to parents of age higher than 10.

All scripts are available at https://github.com/valenzano-lab/NeMAAP-paper to reproduce the results. Detailed model description can be found at https://github.com/valenzano-lab/aegis/wiki/Reference.

## Acknowledgments

We would like to thank all past and current members of the Valenzano lab at the Max Planck Institute for Biology of Ageing and at the Leibniz Institute on Aging for their continuous support and feedback on this project. We are thankful for the support of the Life Science Computing core facility at the Leibniz Institute on Aging, where we ran most of our simulations. Finally, we would like to thank Arian Šajina and William Bradshaw for their significant contribution to earlier versions of AEGIS.

## Authors contributions

M.B. ran all the simulations and generated all data. D.R.V. analyzed the data displayed in Figure 4. D.R.V. conceived the project and wrote the manuscript together with M.B.

## Extended data

**Extended Data Figure 1.**
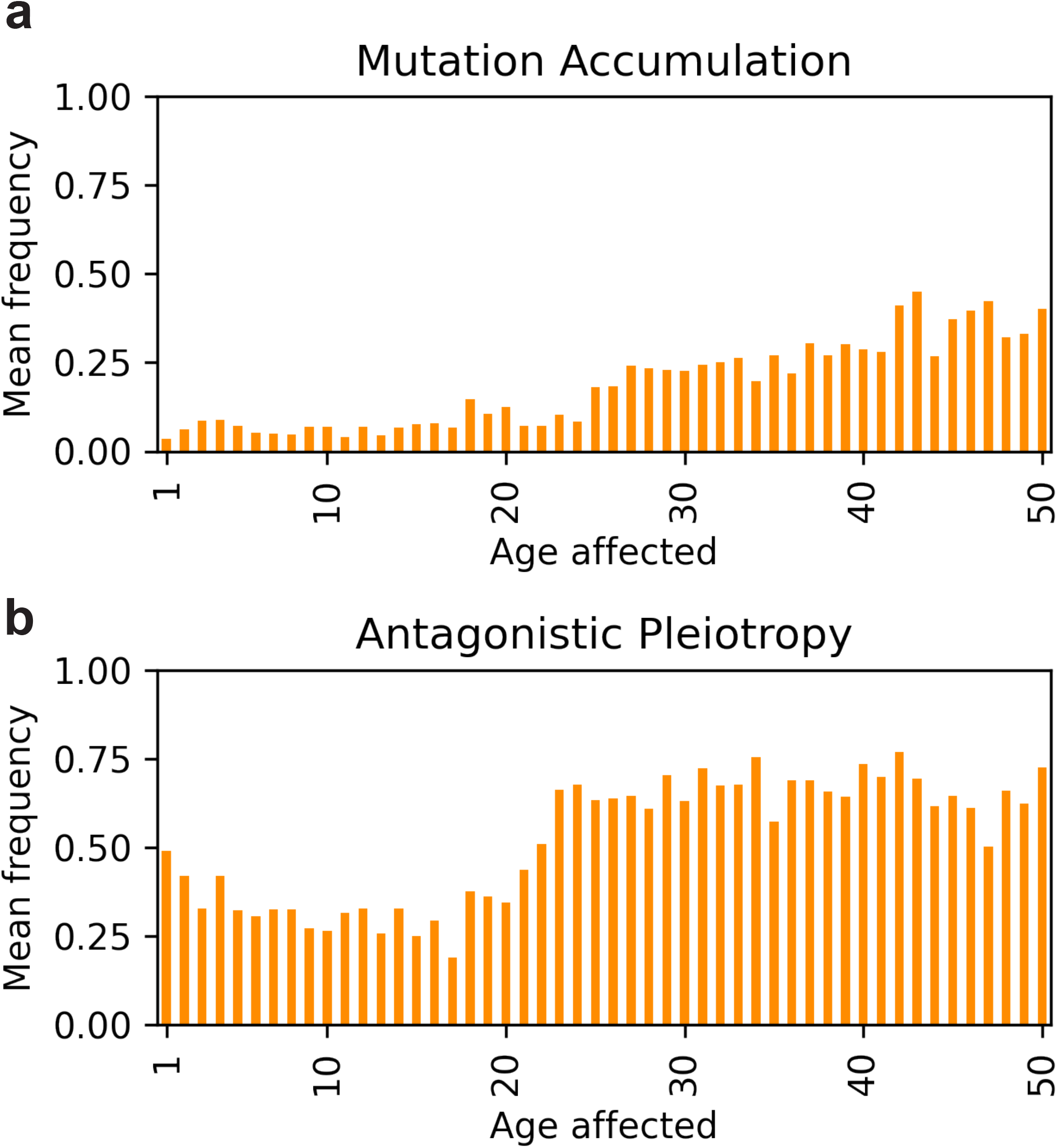
Variants with detrimental effects later in life evolve to higher frequencies when populations evolve under mutation accumulation (a) and antagonistic pleiotropy (b). The y-axis shows the average frequency of variants affecting a specific age (on x-axis).

**Extended Data Figure 2.**
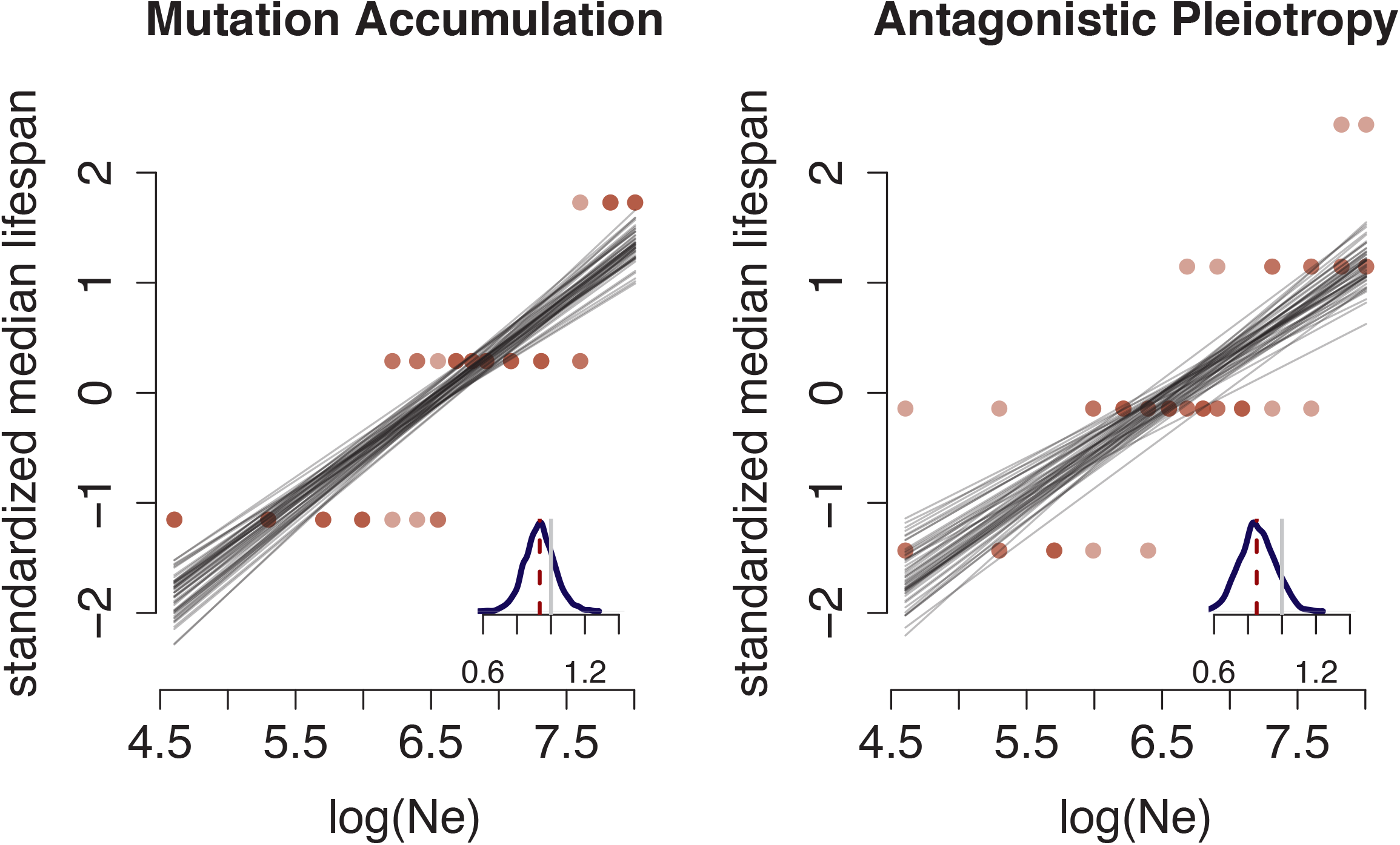
Population size impacts the evolution lifespan. Correlation between standardized median lifespan (y-axis) and population size (log-scale, x-axis) under both mutation accumulation and antagonistic pleiotropy in asexually reproducting populations. Each plot displays 50 regression lines (gray) sampled from the posterior distribution of each regression model. Insets: distribution of regression coefficients (slope); the vertical gray line corresponds to slope = 1 and the vertical dashed red line represents the mean slope for each model.

**Extended Data Figure 3.**
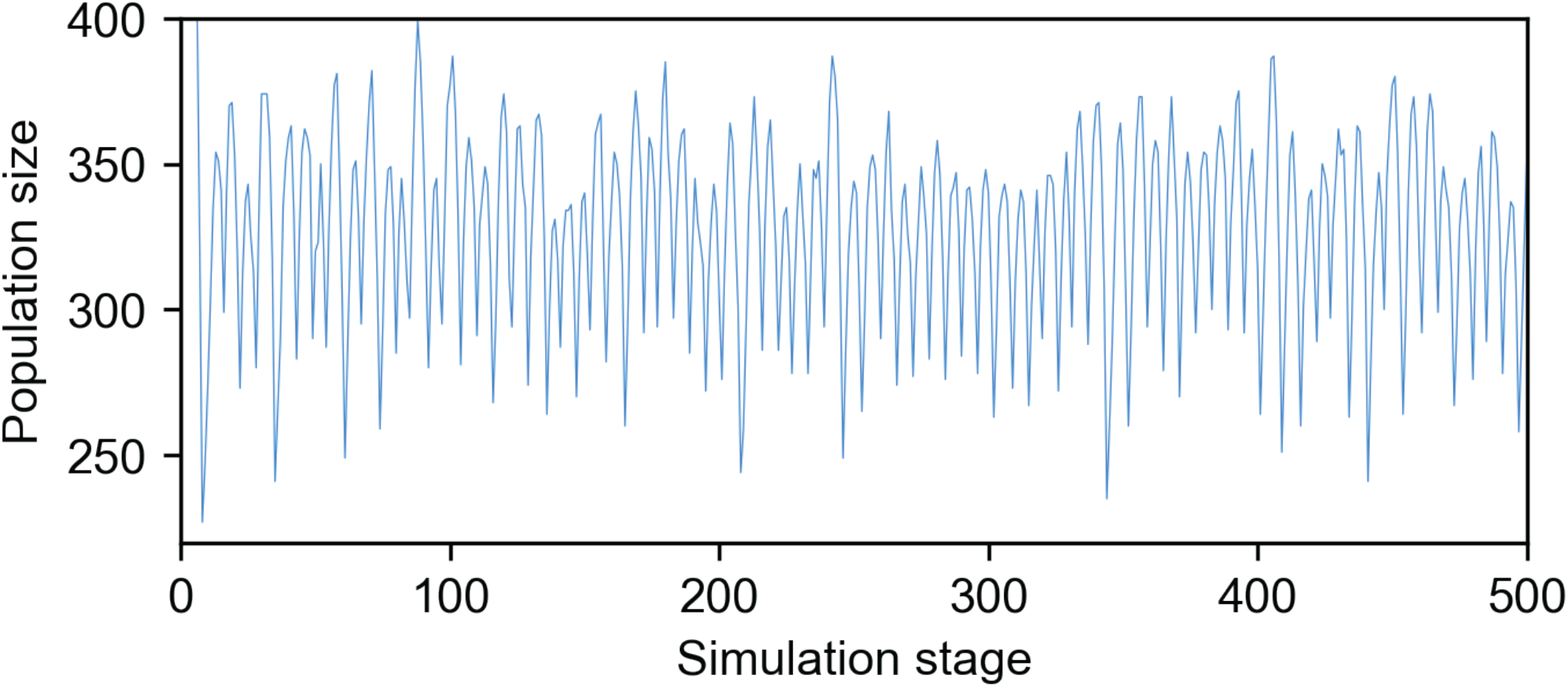
Overpopulation control. When the population overshoots the population size determined by *resource limit*, individuals “starve” resulting in a proportion of the population dying at every stage (**Methods**). The proportion of individuals dying increases with the number of time stages in which the population size exceeds the max allowed population size. In the provided example, the upper allowed population size is set to 300.

